# RNA *post hoc* loading into empty lipid nanoparticles occurs on millisecond timescales during turbulent mixing

**DOI:** 10.64898/2026.06.26.732967

**Authors:** Anas Aljabbari, Heather Binion, Sophia R. Dasaro, Harsa Mitra, Titania Bethiana, Gabriel Harris, Xiaobing Zuo, Mojhdeh Baghbanbashi, Andres Barrio-Zhang, David Perez Herrera, Marxa Figueiredo, Brian K. Wilson, Arezoo Ardekani, Kurt Ristroph

## Abstract

Lipid nanoparticles (LNPs) are conventionally produced through mixing of lipids dissolved in ethanol against a buffer containing RNA. An alternative strategy offering improved cold-chain stability involves formulating empty LNPs (eLNPs), removing ethanol, and *post hoc* loading (PHL) RNA into the aqueous eLNPs. The kinetics of this approach remain unknown. Here, we employ a flowthrough small-angle X-ray scattering (SAXS) setup based on a confined impinging jets (CIJ) mixer to probe PHL kinetics. We show that RNA PHL in a scalable CIJ mixer is efficient and reproducible, and that SAXS data confirms that this process concludes within ∼12 ms under favorable conditions in rapid turbulent micromixing, suggesting a diffusion-limited aggregation mechanism. Favorable conditions were identified as an acidic pH 5.5 buffer combined with turbulent CIJ mixing. In contrast, PHL performed with a neutral pH 7.4 buffer using a CIJ mixer or under laminar flow with a pH 5.5 buffer resulted in inefficient PHL.

## Body

Lipid nanoparticles (LNPs) are multi-component lipid-based nanocarriers that have enabled efficient delivery of fragile nucleic acid therapeutics (siRNA and mRNA), with proven clinical success in the globally distributed COVID-19 vaccines and the gene-silencing drug patisiran^1,2^. The conventional approach to LNP formulation involves rapid and controlled mixing of aqueous acidic buffer containing RNA and ethanol containing lipid components^3^. This leads to co-precipitation of the nucleic acid with the lipids, driven by electrostatic complexation between the polyanion RNA and the charged ionizable lipid component, which forms a structured colloidal dispersion that encapsulates the RNA drug, a process subsequently referred to as *in situ* loading here. While the conventional approach typically yields high (> 90 %) encapsulation and offers protection for the RNA cargo *in vivo*, the precipitated RNA-LNP formulation requires robust and expensive cold chain storage (-80 °C) to maintain formulation integrity^4^. This has posed a major logistical barrier to the broader deployment and distribution of RNA therapeutics, largely due to costs associated with maintaining chemical stability^5^. To address this limitation, emerging research has explored the concept of RNA *post hoc* loading (PHL) into empty LNPs (eLNPs). This strategy involves formulating payload-free (i.e., empty) LNPs by omitting RNA from the aqueous antisolvent buffer in the LNP assembly step and subsequently loading the RNA (in a separate aqueous solution) into the eLNPs, usually through controlling pH, temperature or solvent quality^6–8^. These approaches have yielded successful post-encapsulation of RNA into eLNPs while retaining critical particle physicochemical properties such as particle size, shape, surface charge, and drug loading. Reports show that LNPs loaded through PHL exhibit similar cellular uptake and *in vivo* efficacy when compared to conventional *in situ* loaded LNPs^9^. The enhanced stability conferred by using freeze-dried eLNPs and RNA in this approach further strengthens its applicability as a platform for broader distribution of RNA therapeutics, specifically in regions with limited infrastructure^10^. Importantly, PHL of eLNPs also has the potential to enable highly modular RNA therapeutics, where RNA cargo of choice for cancer immunotherapy, vaccines, or personalized medicine can be incorporated in a plug-and-play manner.

The fundamental mechanistic principles that govern RNA PHL into eLNPs remain underexplored. For both large-scale and point-of-care PHL to be a viable formulation alternative to current *in situ* loading, measuring the kinetics of PHL is essential; understanding the time scale for *post hoc* loading uptake into eLNPs will also be essential in the optimization of the RNA solution and eLNP dispersion mixing technique to minimize compositional heterogeneity among *post hoc* loaded LNPs. Here, we employ our previously-developed flowthrough SAXS platform^11^, in which a confined impinging jets (CIJ) mixer coupled to a quartz capillary flow cell is used to study millisecond-resolution kinetics of processes relevant to nanoparticle self-assembly. Remotely controlled syringe pumps deliver feed streams of eLNPs and RNA into the mixer; by systematically varying the delay tubing length between mixer and capillary, we effectively control fluid residence time at the exposure point, to probe the kinetics of eLNP PHL with RNA at different timepoints downstream of mixing. We also study PHL directly in the mixing chamber using a bespoke engineered CIJ mixer fabricated with a quartz window, which allows SAXS acquisitions at the point of mixing^11^.

### *Post hoc* loading of RNA into eLNPs using a confined impinging jets mixer

Conventional approaches to PHL of RNA into lipid nanoparticles rely on microfluidic or pipette-based mixing of eLNPs and dissolved RNA^8,9^. While these approaches facilitate benchtop academic research, they are limited in their scale-up capacity^12^. To overcome this limitation and assess the feasibility of scalable and continuous PHL, a workflow involving the use of a CIJ mixer (Holland Applied Technologies, IL, USA) was developed (**Figure 1**). Initially, Flash NanoPrecipitation was carried out through mixing of an ethanol (EtOH) stream containing dissolved lipids with an aqueous pH 5.0, 20 mM acetate buffer stream without RNA in a CIJ mixer, to produce a single large batch of eLNPs. Using a syringe pump (PHD Ultra - Harvard Apparatus, MA, USA), the flow rate ratio (FRR) was kept at 1:1 with a total flow rate (TFR) of 120 mL/min (60 mL/min each aqueous and ethanolic streams). The effluent stream containing eLNPs was collected in a pH 5.0, 10 mM acetate buffer quench to lower the EtOH volume fraction to 10% v/v. The resulting eLNP dispersion was dialyzed against a pH 5.0, 10 mM acetate buffer to remove EtOH and subsequently mixed with a separate buffer stream containing RNA (polyA) using a CIJ mixer (detailed formulation parameters in **supporting information**). One batch of eLNPs was used in triplicate for subsequent PHL experiments to ensure identical eLNP particle properties across PHL conditions.

**Figure 1.**
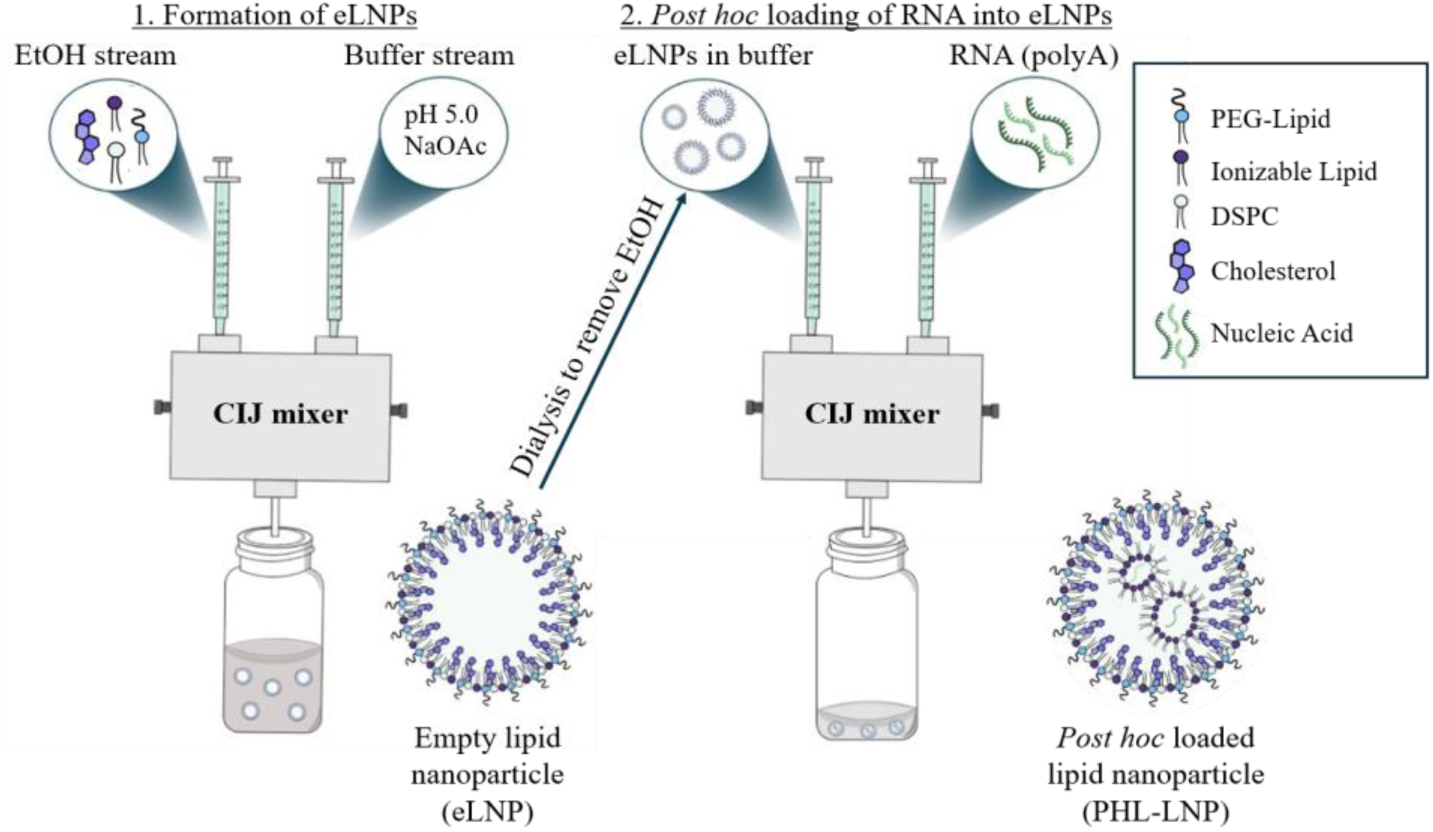
Schematic illustration of two-step PHL workflow for LNPs using a CIJ mixer.

To evaluate the performance of this workflow for the production of PHL LNPs comparable to those reported in the literature, lipid nanoparticles were *post hoc* loaded with polyA (added at an empty LNP ionizable lipid to RNA molar ratio of 5, N/P = 5) under varied pH conditions controlled by a 10 mM acetate or HEPES buffer, ion depending on target pH, in the polyA RNA solution. Dialyzed eLNPs (pH 5.0) were mixed with an aqueous stream of dissolved polyA (variable pH) in a CIJ mixer. Furthermore, to assess the effect of increased outer membrane fluidity on PHL, an additional condition using eLNPs in a pH 4.0, 10 mM acetate buffer containing 25% v/v EtOH was included. Particle size, polydispersity index (PDI), surface charge, and encapsulation efficiency (EE%) were measured for each PHL condition (**Figure 2a-2c**). All PHL conditions were carried out by loading eLNP dispersions and polyA solutions in separate 1 mL plastic syringes, attaching them to the CIJ mixer (**Figure 1**) and rapidly depressing the syringes to achieve turbulent mixing. In agreement with previously reported trends^7^, results show that PHL is highly efficient (EE% > 90 %) at acidic pH, and that efficiency drops significantly in buffer pH values near neutral and above the pKa of the ionizable lipid (EE% < 25 %).

**Figure 2.**
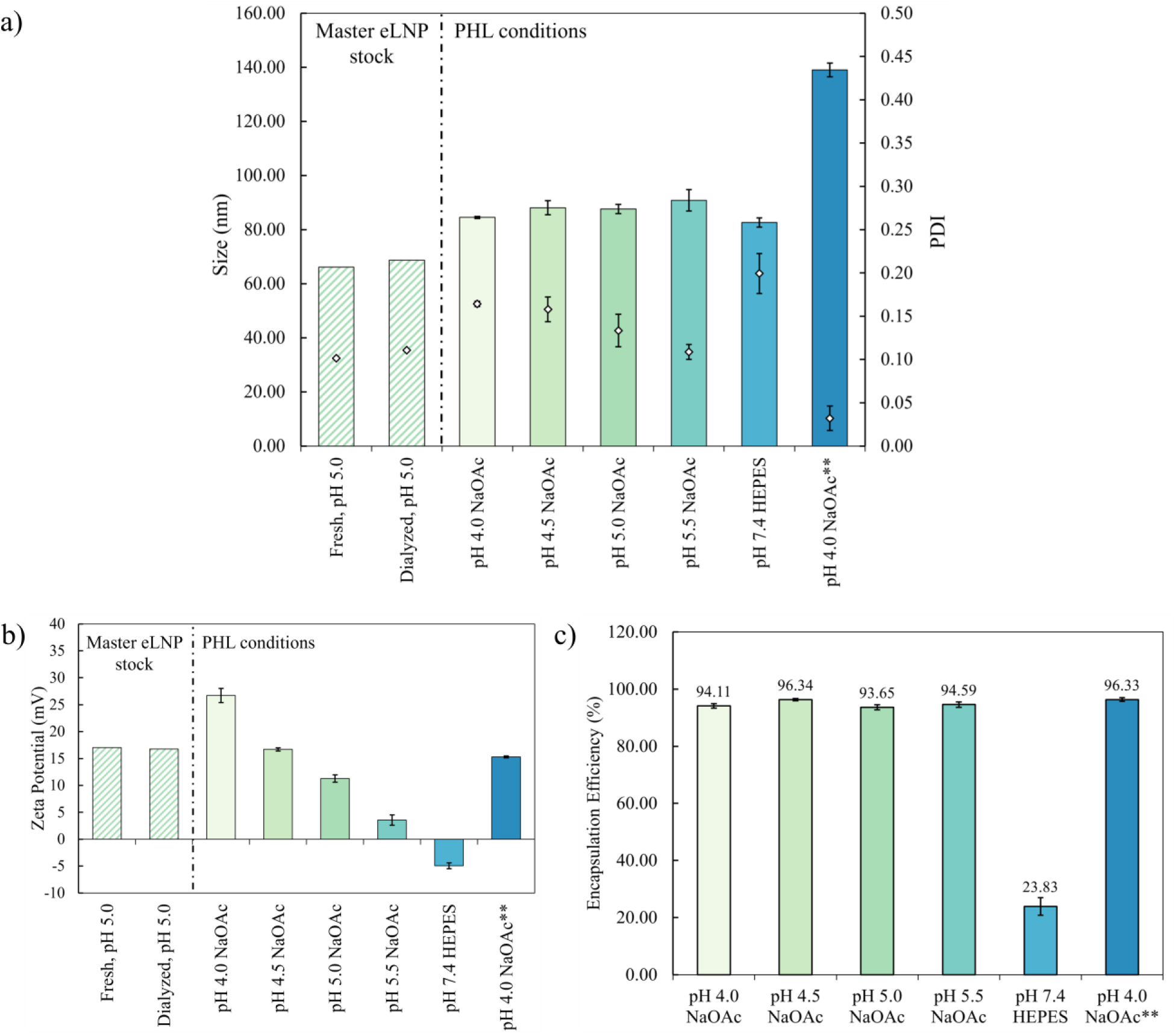
a) Evolution of particle size and PDI due to particle fusion mediated by RNA PHL. Particle size of eLNPs master batch (n = 1) measured pre- and post-dialysis. PHL loading is carried out using eLNPs (in pH 5.0 – acetate buffer) in triplicate with particle size and PDI measured for each replicate (n = 3). Bar graph represents size and points represent PDI. b) Surface charge measurements (zeta-potential) of fresh, dialyzed and PHL LNPs. eLNPs master batch (n = 1) were measured pre- and post-dialysis. PHL loading is carried out using eLNPs in triplicate with zeta-potential measured for each replicate (n = 3). c) Encapsulation efficiency measured for LNPs *post hoc* loaded at lower pH (acidic + acidic w/EtOH) compared to higher pH (neutral). ** In this condition, dialyzed eLNPs are brought to 25% EtOH prior to PHL mixing. Buffer pH values for the eLNP streams and RNA streams, along with the resulting pH in the mixing chamber are summarized in **Table S1**.

As shown in **Figure 2a**, both particle size and PDI increase after mixing with polyA in all tested conditions, other than a drop in PDI observed for PHL at pH 4.0 with 25% EtOH. This is attributed to the adsorption of anionic polyA molecules to the cationic surface of eLNPs, bridging neighboring eLNPs and triggering vesicle fusion, leading to polyA encapsulation in what becomes a larger loaded LNP^6,7^. At pH 7.4, the ionizable lipid is rapidly deprotonated and migrates towards the core of the particle^13,14^, leaving minimal residual surface charge for polyA to interact with for RNA encapsulation. The particle size increase observed at neutral pH is likely to be driven by particle fusion mediated by surface neutralization^15^, rather than nucleic acid bridging, which is also reflected in the low EE% (23.8%). A control experiment was conducted which showed both minimal size increase when mixing eLNPs with RNA-devoid acidic buffers (pH values between 4.0 and 5.5) and larger size increase when mixing eLNPs with a neutral pH 7.4 buffer not containing RNA; results are presented in **Figure S1**, suggesting that the LNP size increase is fundamentally a surface charge neutralization effect, whether due to the buffer pH changing the ionizable lipid charge state or due to partial neutralization and bridging of the cationic surface during RNA adsorption. Interestingly, no significant differences were observed in the EE% of eLNPs PHL in buffer compared to the EE% of eLNPs PHL in the presence of 25% EtOH. This suggests that, under purely aqueous conditions, the outer membrane is sufficiently fluid to allow for efficient particle fusion. It is also noteworthy that in 25% EtOH, *post hoc* loaded LNPs exhibited a significant increase in particle size compared to those in buffer alone (+ 70 nm in EtOH and + 18 nm in aqueous buffer on average). This suggests that increased membrane fluidity and increased solubility of lipid components in the 25% EtOH solvent led to more extensive coalescence, beyond that driven by adsorption of polyA^16^. The surface charges (measured zeta-potentials) of PHL LNPs (**Figure 2b**) scaled as expected, where decreasing the pH of the buffer used in the PHL mixing step increases the cationic charge on the LNP surface. At pH 5.0 and 5.5, a smaller fraction of ionizable lipids is protonated at the surface of eLNPs compared to pH 4.5^17^, which appears to result in a lower surface charge density after adsorption and internalization of polyA. eLNPs *post hoc* loaded at pH 4.0 in aqueous conditions repeatedly displayed a markedly more positive surface charge than initial eLNPs in pH 5.0 buffer, in agreement with the prediction that increasingly acidic pH buffers below the ionizable lipid pKa (ALC-0315 pKa approximately 6.09) charge a larger fraction of the lipid molecules.

In summary, this test showed that turbulent mixing in a CIJ mixer resulted in efficient and reproducible PHL of polyA into eLNPs. These results demonstrate that the PHL formulation workflow using the CIJ mixer is well-suited for reliable flowthrough SAXS analysis.

### Flowthrough SAXS experimental configuration and acquisition parameters

To resolve RNA PHL kinetics at time scales in the tens of milliseconds, continuous flow at high TFR was required. The high TFR ensured (1) good mixing conditions and (2) a high velocity of effluent stream exiting the mixer. Given the dimensions of the outlet tubing and the quartz capillary, the fluid residence time at the point of X-ray exposure can be calculated from the enclosed volume and applied volumetric flowrate (an example of residence time calculations can be found in the **supporting information**). **Figure 3** shows the experimental configuration at the 12-ID-B beamline of the Advanced Photon Source (APS). For acquisitions at the point of mixing, a CIJ mixer with a quartz window was aligned with the X-ray beam path. The CIJ mixer was mounted on a 3D-printed resin holder to secure the mixer during measurements. For acquisitions at longer residence times, polyetheretherketone (PEEK - Cole-Parmer, IL, USA) tubing of varying internal diameters (ID) and lengths was connected between the mixer outlet and the quartz capillary (1.5 mm ID - Charles supper, MA, USA) flow-cell inlet.

**Figure 3.**
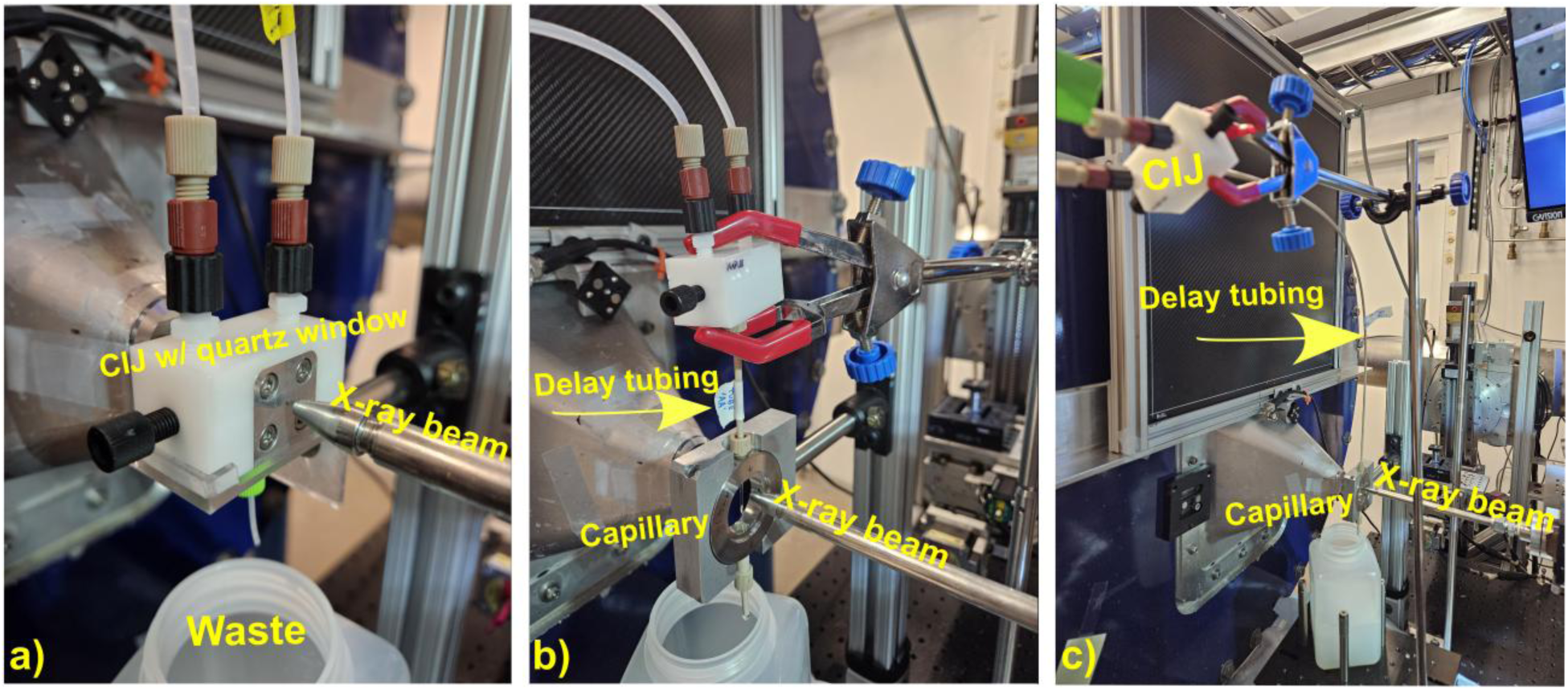
Labelled images of experimental configurations at the 12-ID-B beamline of the Advanced Photon Source. A feed stream of dispersed eLNPs in buffer is mixed with dissolved polyA delivered in the second feed stream at a TFR of 120 mL/min (60 mL/min each stream) a) CIJ mixer with a quartz window (1 mm) aligned with the X-ray beam for frame acquisition at the point of mixing. b) CIJ mixer setup with a delay tubing between the outlet and point of exposure for data acquisition at ∼95 ms fluid residence time. c) CIJ mixer setup with longest delay tubing between the outlet and point of exposure for data acquisition at ∼604 ms fluid residence time.

Remotely controlled syringe pumps were used to deliver two separate streams of eLNPs in buffer and polyA dissolved in buffer at a TFR of 120 mL/min and an FRR of 1:1 (60 mL/min each stream) for all experimental conditions. The mixer was run for a total of 9 s in parallel with SAXS acquisition. Data reduction excluded the first and last 1s of each run to ensure measurement at stable fluid flow (details of acquisition parameters can be found in **supporting information**). Buffer subtraction was carried out using the open-source software BioXTAS RAW^18^.

### Turbulent mixing enables RNA *post hoc* loading into eLNPs within milliseconds under favorable pH conditions

RNA loading into eLNPs produces a distinct SAXS correlation peak that is a result of an internal liquid-crystalline (LC) phase forming in the core of the particle, composed of ionizable lipid aggregates mediated (i.e. templated or nucleated) by the intercalated RNA. By exposing the effluent stream of mixed eLNPs and dissolved RNA to coherent X-rays, the following approach can probe the appearance of an LC phase as a marker of successful PHL^19^. Using the described setup (**Figure 3**), three conditions were tested to determine their respective PHL kinetics. Case 1 was designated as one of the optimal PHL conditions, wherein a stream of pH 5.0 eLNPs was mixed 1:1 with a stream of polyA in pH 5.5 buffer. Case 2 was pH 5.0 eLNPs (brought to 25% EtOH) mixed with polyA in pH 4.0 buffer, to study the effect of increased membrane fluidity on the kinetics of PHL. Case 3 was a control run with eLNPs at pH 5.0 were mixed against polyA in pH 7.4 HEPES buffer (resulting pH: 7.4), a condition where less efficient PHL (EE = 23.8%) was measured. **Figures 4a-d** show the resulting 1D SAXS patterns of each condition at various timepoints after mixing.

**Figure 4.**
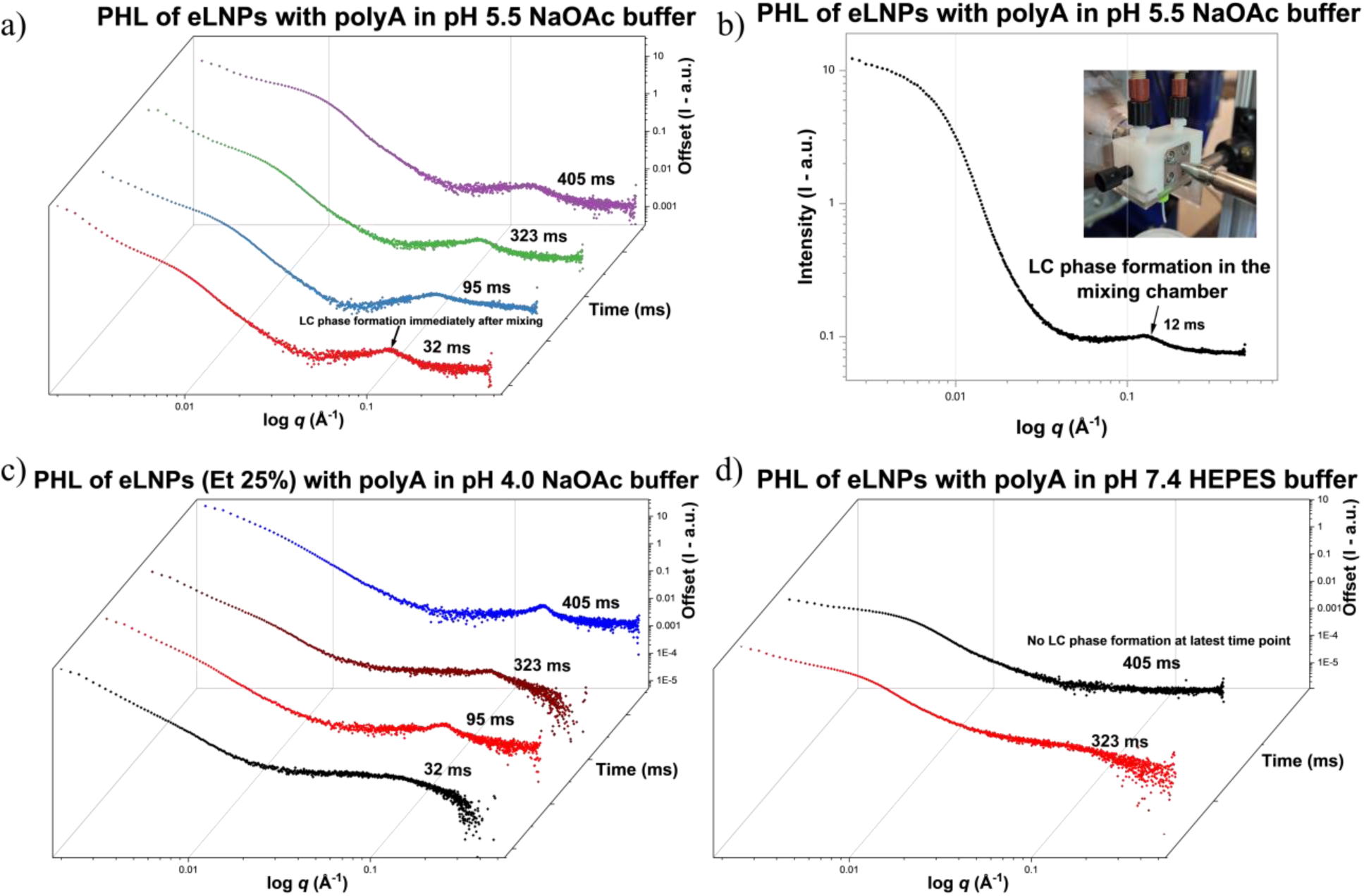
a) Flowthrough SAXS measurements acquired at residence times of 32-604 ms following mixing of pH 5.5 polyA in sodium acetate (NaOAc) buffer and eLNP streams (pH 5.0 acetate buffer). LC phase is observable immediately at earliest timepoint. b) SAXS pattern obtained at pH 5.5 condition at the point of mixing. c) Flowthrough SAXS measurements acquired at residence times of 32-604 ms following mixing of pH 4.0 polyA in NaOAc buffer and eLNP (pH 5.0 NaOAc – brought to 25% EtOH) streams. LC phase formation is delayed and appears at 95 ms after mixing. d) Flowthrough SAXS measurements acquired at residence times of 323-604 ms following mixing of pH 7.4 polyA in HEPES buffer and eLNP streams (pH 5.0 NaOAc buffer). No discernible LC phase formation observed at any measured timepoint.

Results show that the two-step process^6,7^ for PHL appears to be concluded rapidly in the CIJ mixer (**Figure 4a & 4b**) within 12 – 32 ms. The characteristic LC phase that arises due to RNA-ionizable/helper lipid structural rearrangement only appears once RNA is imbibed into the inner core of eLNPs^19^, as opposed to surface association, needing sufficient electron density contrast to be resolved by SAXS (i.e., a sufficient volume threshold of loaded LNPs containing adequate internal volume fraction of nucleated LC phase is required for a detectable diffuse peak in the SAXS plot). Remarkably, a clear LC phase, albeit with lower intensity, is observed even at the point of fluid mixing within the CIJ chamber, confirming the rapid nature of PHL under favorable conditions (**Figure 4b**). It is evident from **Figure 4d** that no LC phase appears even at the latest timepoints measured (604 ms) for PHL with RNA dissolved in neutral pH (7.4 HEPES – 10 mM). While there was still measurable RNA encapsulation (23.8%), the volume of loaded particles did not exceed the threshold for detection by SAXS. This served as a flowthrough benchmark for comparison to PHL at acidic conditions, suggesting that upon appearance of an LC phase, a significant proportion of RNA is already loaded into the core of the eLNPs. It is therefore reasonable to conclude that the majority of eLNPs are loaded by the 12 ms timepoint in **Figure 4b**.

Interestingly, PHL was delayed in the pH 4.0 condition where the eLNPs were spiked to 25% EtOH prior to mixing. The LC phase was not apparent until the 95 ms timepoint. **Figure 2c** shows that final efficiency of PHL with the eLNPs in 25% EtOH is directly comparable to that of PHL in acidic buffers. This suggests that the initial step of PHL, RNA adsorption, may be slowed due to the increased membrane fluidity. The drivers of PHL are the protonated ionizable lipids embedded in the outer lipid layer, which is less ordered in the presence of 25% EtOH in pH 4.0 buffer, resulting in an overall particle irregularity as shown in a previous study at pH 3.0 and 25% EtOH^20^. This likely leads to a delay in favorable electrostatic adsorption at the surface and subsequent particle fusion.

In all cases of efficient PHL, the LC phase that emerged was similar to that which forms with *in situ* loaded LNPs both in the broad peak shape and peak position within 0.125 Å^-1^ – 0.135 Å^-1^ (d-spacing: 46.5 – 50.2 Å). **Figure S2** shows a flowthrough scattering pattern of an *in situ* precipitated RNA-LNP diluted in a CIJ mixer in the same conditions as **Figure 4a**.

### Laminar fluid mixing is associated with slower RNA *post hoc* loading

Next, we tested how mixing efficiency would affect the kinetics of PHL. A Precigenome MIX-4 (Precigenome - CA, USA), a tesla micromixer chip, was used to achieve laminar flow fluid mixing. **Figure S3** shows an image of the experimental setup at the beamline. At an FRR of 1:1 and a TFR of 4 mL/min, addition of tubing between the outlet of the micromixer and the quartz capillary resulted in a significant increase in fluid residence time compared to the CIJ setup, due to the low flow rate through the MIX-4 mixer. The setup shown in **Figure S3** resulted in a fluid residence time of ∼1500 ms at the point of exposure, an order of magnitude higher than the fastest timepoints resolved using the high TFR in the CIJ mixer.

Figure 5c shows the scattering pattern of LNPs *post hoc* loaded using a micromixer at optimal pH conditions, with a fluid residence time of ∼1500 ms. The LC phase peak observed at this time point was broader and weaker in intensity than those observed in Figure 4. While the low-*q* overall particle scattering of the empty eLNPs was clearly detected, it was evident that a significant proportion of them were yet to be loaded with RNA, even at ∼1500 ms after mixing. In contrast to the results observed using a CIJ mixer, the majority of PHL does not seem to occur in the micromixer itself but downstream in the collection vessel of the effluent fluid. This suggests that although PHL in a micromixer ultimately yields high RNA encapsulation, this process likely occurs over significantly longer timescales, dictated by the diffusivity of the eLNPs and the RNA in solution. Consistent with our observations, prior work using Förster resonance energy transfer (FRET) spectroscopy to probe LNP fusion kinetics during gradual neutralization (low flowrate) reported population level particle fusion timescales of tens to hundreds of seconds^21^.

**Figure 5.**
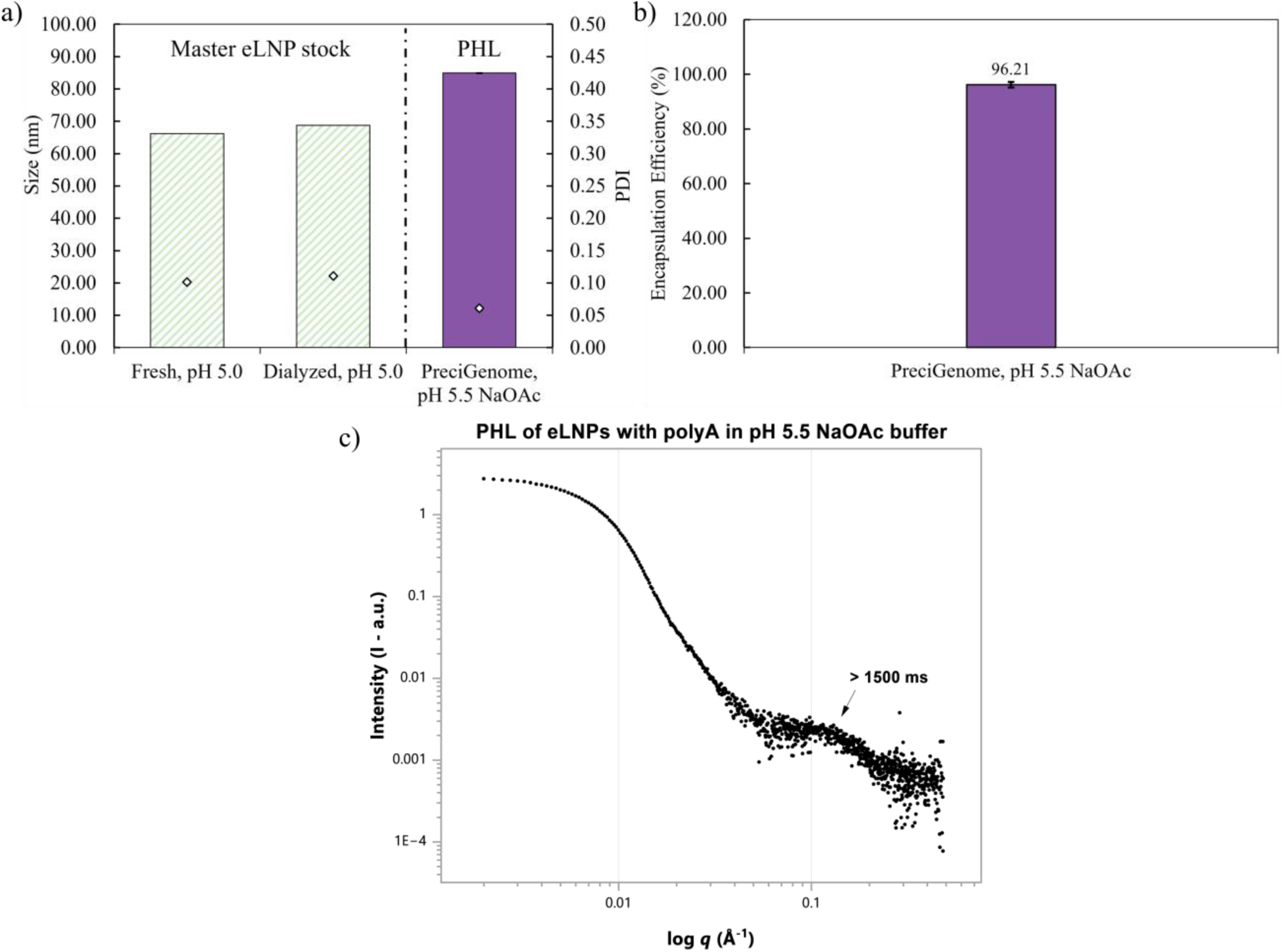
a) Results for size and PDI for particles *post hoc* loaded with polyA in pH 5.5 NaOAc buffer using a MIX-4 micromixer. Particle size and PDI of eLNPs master batch (n = 1) measured pre- and post-dialysis. PHL loading is carried out using eLNPs in triplicate with particle size and PDI measured for each replicate (n = 3). Bar graph represents size and points represent PDI. b) Encapsulation efficiency measured for LNPs *post hoc* loaded with polyA at pH 5.5 in a micromixer – this measurement took place more than 10 minutes after formulation and does not refer to the encapsulation at the ∼1500ms time point. c) Flowthrough SAXS measurements acquired at a residence time of ∼1500 ms following mixing of pH 5.5 polyA in sodium acetate (NaOAc) buffer and eLNP streams (pH 5.0 NaOAc buffer) in a micromixer. LC phase peak observed is broad with low intensity, suggesting incomplete PHL.

### Timescales of *post hoc* loading are governed by diffusion length scales

The timescale over which PHL of nucleic acids into eLNPs occurs is governed by diffusion over characteristic distances between discrete eLNPs and hydrodynamically large, solvated RNA polymer coils. Mixing alters these effective length scales by stretching and folding fluid lamellae of each inlet stream, which also acts to blend the concentration fields, thereby accelerating homogenization by reducing the average inter-species distance an RNA coil must diffuse before adsorbing to an eLNP. The extent of this enhancement depends on mixer geometry and flow conditions. Understanding the diffusion-limited kinetics of this process is therefore essential for rationalizing PHL efficiency under the two different mixing regimes examined herein. Turbulent mixing in the CIJ utilizes the energy dissipating vortex structure of the flow field to generate roughly 1 micron thick lamellae of each inlet fluid with a mixing timescale on the order of 1 ms^22^ while laminar flow in the microfluidic device relies on the physical dimensions of the flow channel to produce lamellae on the order of 100 microns thick with a mixing timescale on the order of 10^2^ –10^3^ of ms^23^.

A previous study has applied a diffusion timescale model to show that homogenous encapsulation of hydrophobic iron oxide colloids (varying sizes) and hydrophobic polystyrene into nanoparticles stabilized with amphiphilic polystyrene-*b*-PEG depends on matching the diffusivity timescales of the two hydrophobic components^24^. The model, supported by experimental data, revealed a critical threshold of relative diffusivities that must be satisfied to achieve stoichiometric encapsulation of differently-sized components within a single nanocarrier. We employ a similar framework to analyze PHL of polyA RNA into eLNPs, using an average diffusion timescale of the participating components (fully formed eLNPs and dissolved RNA) rather than a ratio of individual diffusivities. This approach enables direct comparison between diffusion-limited estimates in each regime and the observed PHL kinetics measured by flowthrough SAXS.

Eqn. 1 defines the characteristic diffusion timescale (τ_i_), which is a function of the distance traversed (*l*) and the diffusivity, *D*_i_. The diffusivity is determined using the Stokes-Einstein equation (Eqn. 2), where *k_B_* is Boltzmann’s constant, *T* is the absolute temperature, *η* is the solvent viscosity, and *R_i_* is the hydrodynamic radius of species *i*.

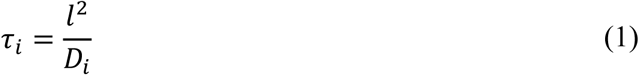

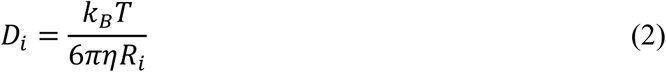

Under homogeneous, well-mixed conditions, the characteristic diffusion length required for an eLNP to encounter an RNA molecule is given by the mean surface-to-surface separation, expressed in Eqn. 3. This separation depends on the particle volume fraction, *φ*_i_, and the maximum close-packed volume fraction for spheres, *φ*_max_ = 0.64. For RNA and eLNPs alike, the volume fraction is determined directly from the RNA mass concentration *C*, normalized by its density *ρ.* Accordingly, Eqn. 4 accounts for the mass concentration and density in the volume fraction of each diffusing species.

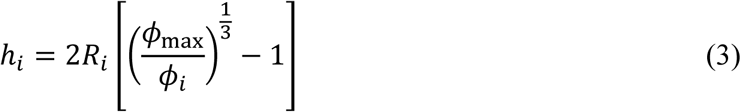

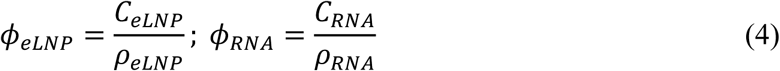

Combining these expressions, an average diffusion timescale can be calculated using Eqn. 5, in which the surface-to-surface separation and diffusivity of eLNPs and polyA RNA across the relevant length scales are number-averaged **(supporting information)**. This represents the time required for an average RNA molecule to diffuse to the surface of an eLNP. RNA adsorption to the eLNP surface is assumed to be highly efficient (all RNA-eLNP collisions are productive at the optimal PHL pH) due to the multivalent electrostatic interaction between the cationic eLNP surface and RNA polyanion.

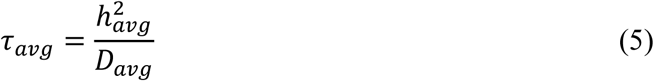

For polyA RNA lengths which range from 300-4400 nucleotides in the material used in this study, turbulent mixing yields an average diffusion timescale for the components of less than 10^-2^ s (**Figure S4 & S5**)^22^. This estimate is in close agreement with the experimentally observed PHL timescale of approximately 12 ms measured in the CIJ mixer using flowthrough SAXS, under acidic conditions. SAXS profiles show modest evolution after this 12 ms time point, suggesting that additional RNA loads into the LNP and furthers internal LC formation but most of the RNA is captured by this theoretically calculated 10 ms timescale for the average RNA molecules to diffuse to (and rapidly imbibe into the core of) an eLNP. The increased viscosity of a 25% ethanol solutions (η_s_ about 2.3 cP) also contributes significantly to the delayed PHL-indicating LC phase SAXS peak evolution in Figure 4c by doubling the time for RNA to diffuse to an eLNP.^25^ LC peak evolution between 32 ms and 95 ms in this ethanol-containing case strongly hints that the fundamental aqueous PHL timescale is around the 15-30 ms timescale measured from the in-mixer data.

In contrast, transitioning from turbulent, homogeneous mixing to laminar flow conditions substantially alters the characteristic length scales over which diffusion must occur. Under laminar flow, where eLNPs and RNA are assumed to reside in interfacing aqueous lamellae, productive collisions require diffusion across much larger distances prior to PHL. As illustrated in Figure 6, the characteristic diffusion length under laminar flow is governed by the channel width of the mixer rather than interparticle separation. Assuming an average position at the center of each lamella, the diffusion length becomes half the channel width. Using a representative microfluidic channel width of 0.1 mm, applying Eqn. 1 yields average diffusion timescales on the order of 2 x 10^2^ s for both eLNPs and RNA. Despite this prolonged timescale of diffusion, these results do not imply that PHL is entirely absent at shorter times but rather indicates that homogeneous loading across the particle population is significantly delayed relative to turbulent mixing.

**Figure 6.**
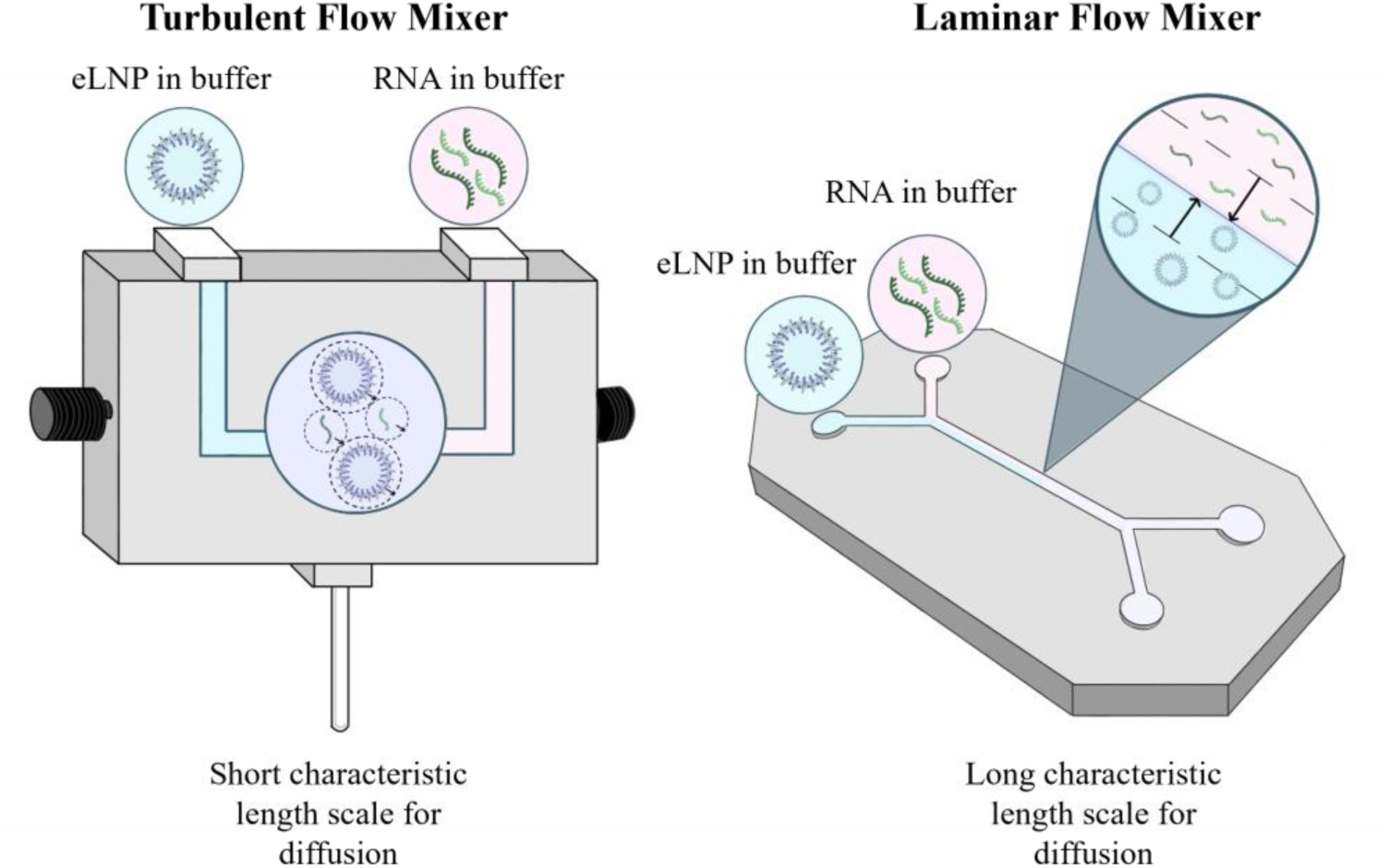
Illustration of two mixing regimes analyzed in the PHL workflow. The characteristic length scale of diffusion for eLNPs and RNA for turbulent flow mixing depends on the surface-to-surface separation between eLNPs and RNA. For laminar flow mixing, the characteristic length scale is a function of the channel width.

These predictions are consistent with the SAXS-derived PHL kinetics shown in Figure 5c, where a weak LC phase peak emerges after a fluid residence time of approximately 1500 ms. Since the LC phase signal scales with the volume fraction of nanoparticles exhibiting an internal structure, its subtle appearance indicates an LNP population partially (and incompletely) loaded with polyA RNA complexed in the core.

In summary, we have shown using a bespoke flowthrough SAXS setup, that RNA PHL into eLNPs using turbulent mixing in an acidic pH regime concludes rapidly within 12-32 ms, suggesting a self-assembly mechanism that is diffusion-limited. The efficiency and kinetics of this process are significantly hindered in neutral buffer pH and by substituting turbulent mixing with microfluidic mixing. This highlights that homogeneous PHL likely requires optimal conditions in the form of fluid mixing faster than the time scale for diffusion-limited aggregation, as well as a favorable acidic pH environment. Together, these findings inform both the design of industrial PHL manufacturing strategies, as well as point-of-care production of RNA-loaded LNP therapeutics.

## Acknowledgements

This research was performed on APS beam time awards (DOI: https://doi.org/10.46936/APS-189691/60013831 & https://doi.org/10.46936/APS-191740/60015470) from the Advanced Photon Source, a U.S. Department of Energy (DOE) Office of Science user facility operated for the DOE Office of Science by Argonne National Laboratory under Contract No. DE-AC02-06CH11357. This research was supported by a grant from Eli Lilly and Company (USA), part of the Eli Lilly and Purdue University Research Alliance Center (LPRC). We gratefully acknowledge Lucas D. Johnson for fruitful discussions on the diffusivity timescales modelling.

