## Supporting information for "RNA *post hoc* loading into empty lipid nanoparticles occurs on millisecond timescales during turbulent mixing"

### Methods section

#### Materials:

ALC-0315 (6-((2-hexyldecanoyl)oxy)-N-(6-((2-hexyldecanoyl)oxy)hexyl)-N-(4-hydroxybutyl)hexan-1-aminium), DSPC (1,2-distearoyl-sn-glycero-3-phosphocholine), cholesterol, and DMG-PEG2000 (1,2-dimyristoyl-rac-glycero-3-methoxypolyethylene glycol-2000) were purchased from Avanti Polar Lipids (Alabama, USA). Polyadenylic acid potassium salt (polyA) was purchased from Sigma Aldrich (Missouri, USA). Absolute ethanol (99.8%) and buffer salts (HEPES free acid, sodium acetate) were purchased from Fisher Scientific (Massachusetts, USA). Quant-iT Ribogreen® RNA assay kit and Triton® X-100 were purchased from ThermoFisher Scientific (Massachusetts, USA).

#### Lipid nanoparticle formulation and process parameters

RNA-LNPs and eLNPs were produced by Flash NanoPrecipitation achieved through mixing of an ethanolic stream containing dissolved lipids and an aqueous stream containing 20 mM sodium acetate solution (pH 5.0) either with or without dissolved polyA (0.6 mg/mL), respectively. The lipid stream contained a 12 mg/mL mixture of ALC-0315, DSPC, cholesterol and DMG-PEG2000 at a molar ratio of 50/10/38.5/1.5 mol%, respectively. Mixing was carried out in a CIJ mixer using syringe pumps (PHD Ultra - Harvard Apparatus, MA, USA) maintaining a total flow rate (TFR) of 120 mL/min and a flow rate ratio (FRR) of 1:1. The effluent stream was immediately quenched using 10 mM sodium acetate buffer (pH 5.0) to bring the ethanol volume fraction down to 10% v/v, resulting in a total lipid concentration of 1.2 mg/mL. The resulting RNA-LNPs/eLNPs were dialyzed against 10 mM pH 5.0 sodium acetate buffer to remove ethanol. Dialysis was carried out at RT in a Spectra/Por® 1 dialysis membrane (MWCO of 6-8 kDa) against a buffer volume equivalent to 50x the RNA-LNP/eLNP suspension volume for a duration of 3 hours. To ensure adequate SAXS data quality and resolution, dialyzed eLNPs/LNPs for SAXS analysis were concentrated (5x) using Amicon centrifugal filters (MWCO 100 kDa, Millipore Sigma) to achieve a total lipid concentration of 6 mg/mL. Filters were primed at 500 x *g* for 2 min with a 70:30 (v/v) RNase-free water:EtOH mixture, followed by two water-only steps. Centrifugation was done in an Eppendorf 5430 centrifuge (Hamburg, Germany) at 500 x *g* (centrifugation time = ~ 4 min per mL of filtrate). Dialyzed and concentrated eLNPs/LNPs were stored at 4 °C until further use.

### *Post hoc* loading of polyA into eLNPs

*Initial lab-based CIJ mixer PHL experiments:*

Using a CIJ mixer, dialyzed eLNPs with a total lipid concentration of 1.2 mg/mL were mixed with a buffer stream containing 0.06 mg/mL polyA (N/P: 5) in either 20 mM sodium acetate buffer or 20 mM HEPES buffer, over a pH range of 4.0 – 7.4 (4.0, 4.5, 5.0, 5.5 and 7.4). This also included a condition where eLNPs were brought to 25% EtOH and mixed with polyA in a pH 4.0 acetate buffer. Each stream was loaded into a 1 mL plastic syringe (0.55 mL per stream) and attached to the CIJ inlets. Turbulent mixing was achieved by rapidly depressing the syringes and collecting the effluent stream of *post hoc* loaded LNPs in a scintillation vial.

*Initial lab-based Precigenome micromixer PHL experiment:*

Using a Precigenome MIX-4 micromixer, dialyzed eLNPs with a total lipid concentration of 1.2 mg/mL were mixed with a buffer stream containing 0.06 mg/mL polyA (N/P: 5) in 20 mM sodium acetate buffer. Each stream was loaded into 3 mL plastic syringes and attached to the micromixer inlets. Laminar flow mixing at a TFR of 4 mL/min and a FRR of 1:1 was achieved using syringe pumps. Effluent stream of *post hoc* loaded particles was collected in a scintillation vial.

*Synchrotron small angle X-ray scattering (SAXS) studies:*

For PHL studies carried out at Argonne National Laboratory, pre-made eLNPs and RNA-LNPs (dialyzed and 5x concentrated to 6 mg/mL) were transported to the beamline at 4 °C and were used within 72 hours of initial preparation. PHL was carried out in a CIJ mixer at a TFR of 120 mL/min at an FRR of 1:1. In practice, 60 mL/min eLNP stream and 60 mL/min buffer stream containing 0.3 mg/mL polyA. TFR and FRR were set to 4 mL/min and 1:1 for eLNPs loaded with polyA using the Precigenome micromixer. All analyzed samples had a total lipid concentration of 3 mg/mL and a total polyA concentration of 0.15 mg/mL at the point of X-ray exposure.

### Physicochemical characterization of eLNPs and RNA-LNPs

Particle size distributions (hydrodynamic diameter), PDI and surface charge (zeta-potential) were determined using dynamic light scattering (size, PDI) and electrophoretic light scattering (zeta-potential), respectively, on a Malvern Zetasizer Pro (Malvern Panalytical, United Kingdom), at a

173° scattering angle. Measurements were performed in triplicate, and all samples were diluted 10-fold in buffer prior to analysis.

RNA encapsulation efficiency (EE%) was determined using a Quant-iT Ribogreen® RNA assay kit. Briefly, RNA content encapsulated in LNPs was determined by comparing fluorescence signals measured in the presence and absence of 0.5% Triton X-100 (lysing detergent) in Tris-EDTA buffer. Fluorescence measurements were carried out on a Molecular Devices SpectraMax i3x (CA, USA) using an $\lambda_{excitation}$ = 485 nm and $\lambda_{emission}$ = 528 nm as recommended by the manufacturer for the Ribogreen® dye. EE% was calculated by comparing total RNA in the sample to the fraction of unencapsulated RNA using the following equation:

$$EE\% = \frac{Total\ RNA - Unencapsulated\ RNA}{Total\ RNA}\ x\ 100$$

**Flowthrough synchrotron small-angle X-ray scattering**

Flowthrough SAXS experiments were carried out in air in transmission mode at the 12-ID-B beamline of the Advanced Photon Source, Argonne National Laboratory (IL, USA). Samples were flowed through a thin-walled quartz capillary with a 1.5 mm diameter and a wall thickness of 10 microns (Charles supper, MA, USA) in an in-house fabricated flow-cell. Frames were acquired using a fixed X-ray energy of 13.3 KeV (λ = 0.932 Å) and an Eiger2 9 M X-ray detector (DECTRIS). The sample to detector distance (SDD) was set to 3.6 m with silver behenate (AgBeh) as calibration standard to determine the $q$-scale (~0.002 – 0.48 $Å^{-1}$) where $q$ denotes the scattering vector, defined as:

$$q = \frac{4\pi}{\lambda}\sin(\theta)$$

where λ is the X-ray wavelength and θ is the scattering angle. The $q$ positions of scattering peaks were used to determine the d-spacings of liquid crystalline domains (internal ordering) in the lipid nanoparticles, using the following relationship:

$$d = \frac{2\pi}{q}$$

In all flowthrough experiments, X-ray exposure time was set to 50 ms with a total of 150 frames per acquisition (9 seconds of fluid flow) and a 10 ms delay between each frame acquisition. The initial 5 frames and the final 45 frames were omitted from preprocessing and subsequent data analysis, as these intervals were designated as pre- and post-waste (i.e., periods of unstable fluid flow). Background scattering patterns with the relevant background compositions were collected before the start of each sample run. In between runs, the mixer and the capillary were flushed with water and EtOH for cleaning.

**Example of fluid residence time calculation:**

For delay tubing that gives SAXS acquisitions at ~604 ms, PEEK tubing with internal diameter (ID) of 0.00158 m and a length of 0.6 m (L) was used. The ID of the quartz capillary was 0.0015 m (ID), and the X-ray beam was positioned at 0.005 m (L) from the top, which was the length of the capillary that the fluid travels through before exposure.

$$Volume\ (604\ ms\ setup) = \frac{\pi\ (ID)^2}{4} * L \tag{1}$$

$$Volume\ (604\ ms\ setup) = \frac{\pi\ (0.00158)^2}{4} * 0.6\ = 3.14\ x\ 10^{-08}\ m^3 \tag{2}$$

Flow rate (Q) expressed in 0.000002 $m^3/s$ was determined from the volumetric flow rate of 120 mL/min. Residence time in CIJ mixer ($t_{mixer}$) at 120 mL/min was determined to be 12 ms. Residence time in 0.005 m of quartz capillary ($t_{capillary}$) at 120 mL/min was determined to be 4.4 ms.

$$Residence\ time\ (t_{total}) = \left(\frac{V}{Q}\right) + t_{mixer} + t_{capillary} \tag{3}$$

$$Residence\ time\ (t) = \left(\frac{3.14\ x\ 10^{-08}\ m^3}{0.000002\ {m^3}/{s}} * 1000\right) + 12\ + 4\ = 604.3\ ms \tag{4}$$

**Table S1.** Buffer conditions of the two inlet streams. The pH in the mixing chamber was estimated by mixing equal volumes of the inlet streams and measuring the pH of the resulting mixture using a pH probe.

| pH - eLNP stream<br>Ionic strength: 10 mM | pH - RNA stream<br>Ionic strength: 10 mM | pH in mixing chamber (CIJ) |
|---|---|---|
| 5.0 | 4.0 | 4.65 |
| 5.0 | 4.5 | 4.88 |
| 5.0 | 5.0 | 5.0 |
| 5.0 | 5.5 | 5.30 |
| 5.0 | 7.4 | 7.4 |

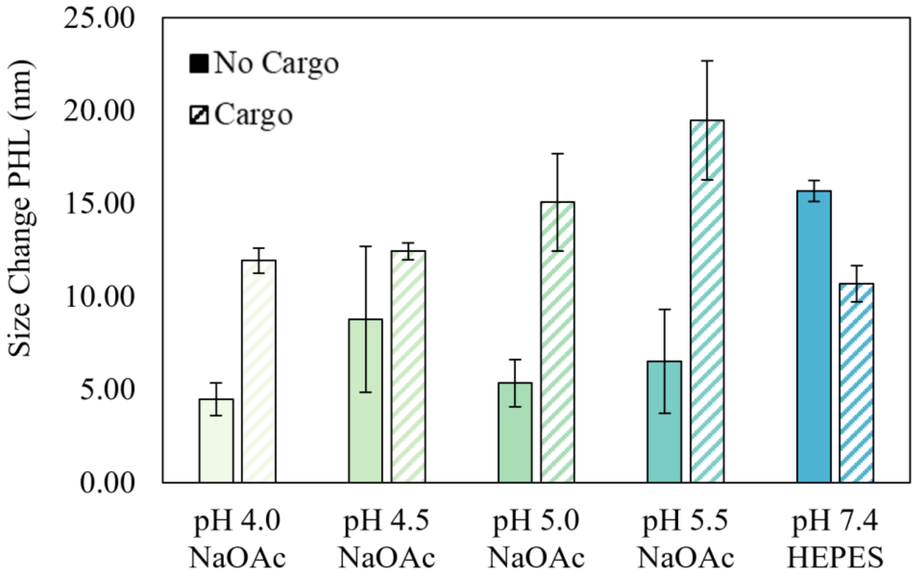


**Figure S1.** Comparison of particle size changes following mixing in a CIJ mixer of eLNPs with buffer streams containing (dashed bars) or lacking (filled bars) RNA cargo. Size change is significantly larger when eLNPs are mixed with buffer containing RNA.

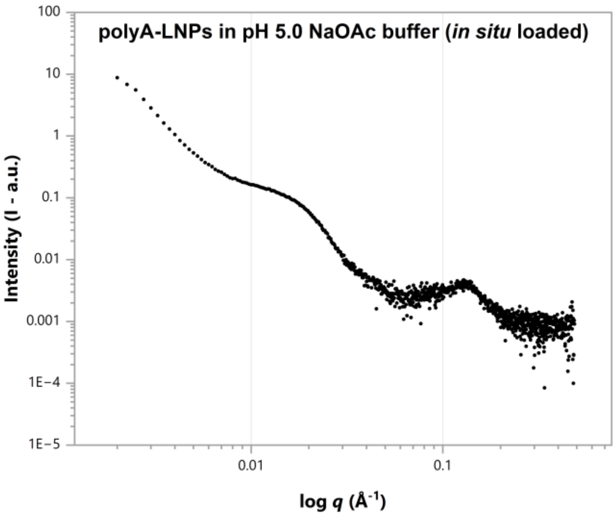


**Figure S2.** Flowthrough SAXS pattern of *in situ* precipitated and dialyzed RNA-LNP in pH 5.0 sodium acetate buffer, following mixing with pH 5.5 sodium acetate buffer. Peak shape and position serve as benchmark for comparison to LC-phase peak observed in PHL.

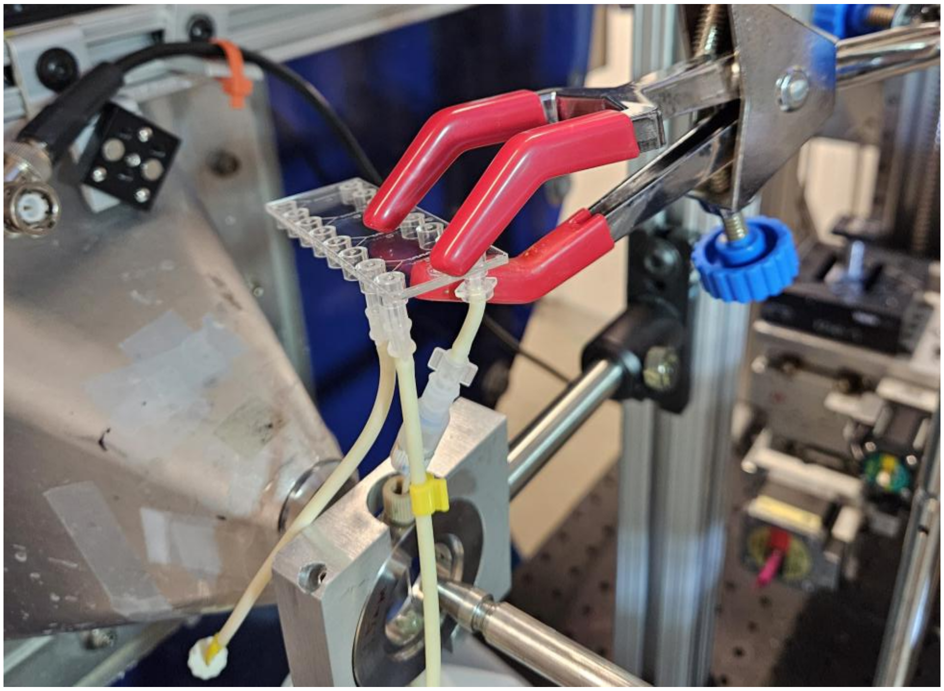

**Figure S3.** Image of micromixer experimental configuration at the 12-ID-B beamline. Flow from the outlet of the MIX-4 chip is directed into the quartz capillary flow cell.

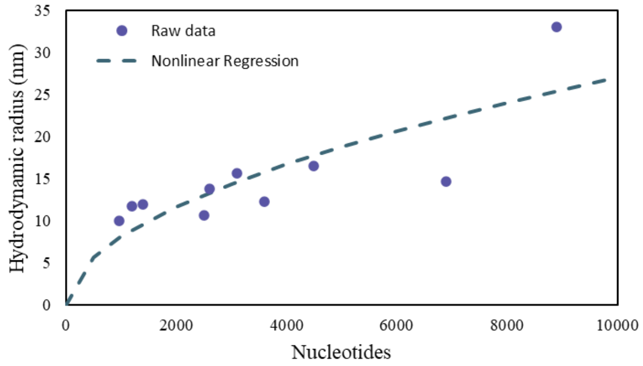


$$Hydrodynamic\ radius\ (nm) = 0.218\ x\ no.of\ nucleotides^{0.523}$$

**Figure S4.** Single stranded RNA data correlating number of nucleotides to hydrodynamic radius (nm) using nonlinear regression methods on MATLAB[1,2].

$$D_{avg} = \frac{\phi_{eLNP} D_{eLNP} + \phi_{RNA} D_{RNA}}{\phi_{Total}} \quad (1)$$

$$h_{avg} = \frac{h_{eLNP} + h_{RNA}}{2} \quad (2)$$

Number averaged diffusivity ($D_{avg}$) of eLNPs and RNA are calculated as a function of volume packing parameters. Here, $\phi_{Total}$ represents the combined volume fractions of eLNPs and RNA. Averaged surface-to-surface separation distances ($h_{avg}$) are calculated for this dual system.

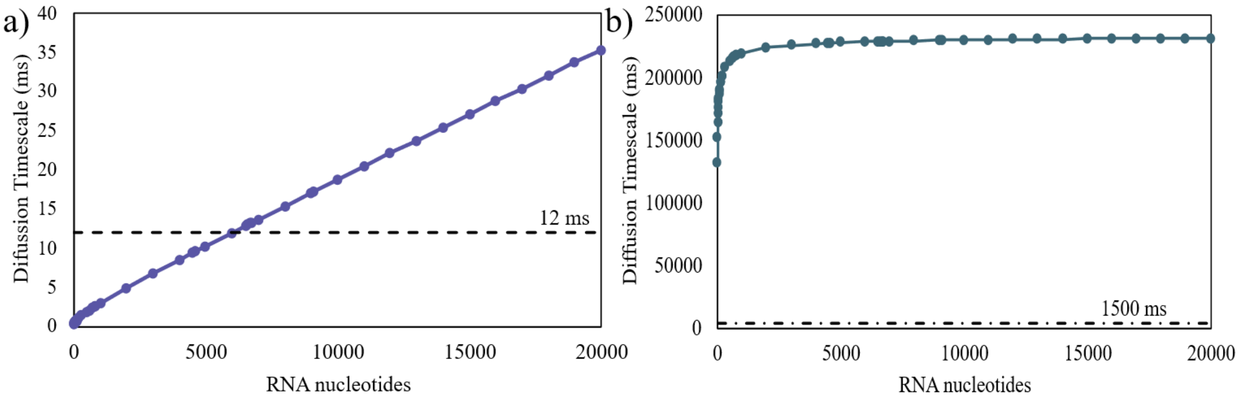


**Figure S5.** Diffusion time scale, and consequently PHL kinetics, as a function of RNA size for a) turbulent flow mixing with Kolmogorov length scale of diffusion and b) laminar flow mixing with macroscopic channel width-based length scale of diffusion.

### <u>References</u>

(1) Borodavka, A.; Singaram, S. W.; Stockley, P. G.; Gelbart, W. M.; Ben-Shaul, A.; Tuma, R. Sizes of Long RNA Molecules Are Determined by the Branching Patterns of Their Secondary Structures. *Biophys. J.* **2016**, *111* (10), 2077–2085. https://doi.org/10.1016/j.bpj.2016.10.014.

(2) The MathWorks, Inc. MATLAB.
